# DNA methylation signatures of mismatch repair-deficient colorectal cancer

**DOI:** 10.64898/2026.04.09.717165

**Authors:** Rebecca Ward, Molly Endicott, Bethan Mallabar-Rimmer, Joe Burrage, Kitty Sherwood, Qiwen Huang, Joseph C. Ward, Steve Thorn, Connor Woolley, Sophie J. Wood, Emma Dempster, Harry D. Green, Ian Tomlinson, Amy P. Webster

## Abstract

**Background:** Colorectal cancer (CRC) is a molecularly heterogeneous disease shaped by both genetic and epigenetic alterations. Approximately 15% of CRCs display widespread CpG island hypermethylation, known as the CpG Island Methylator Phenotype (CIMP). CIMP-high (CIMP-H) tumours frequently exhibit *MLH1* promoter hypermethylation, leading to mismatch repair deficiency (MMRd) and microsatellite instability (MSI). However, DNA methylation patterns associated with MSI, independent of CIMP and *MLH1* silencing, and the influence of clinical variables such as anatomical location and patient age on the CRC methylome remain poorly characterised.

**Methods:** We performed epigenome-wide DNA methylation profiling of 259 primary CRC tissue samples using the Illumina EPICv2 array, comparing differential methylation between MSI and microsatellite stable (MSS) CRC, adjusting for tumour purity, *MLH1* promoter methylation, CIMP status, and anatomical location, to account for known confounders. We further evaluated the independent effects of anatomical location and patient age on global methylation patterns.

**Results:** Epigenome-wide differential methylation between MSS and MSI CRC was dominated by *MLH1* promoter hypermethylation. After adjusting for *MLH1* hypermethylation and CIMP status, we identified a distinct set of 656 CpG sites associated with MMRd independent of *MLH1* silencing. These included hypermethylation at *LRP6*, *GSK3β,* and *CDK12,* implicating altered WNT signalling and transcriptional regulation pathways. Comparison of MSI subgroups revealed the co-occurrence of *MLH1* hypermethylation with promoter hypermethylation at *TXNRD1*. Anatomical location showed a strong independent effect on methylation patterns, while we observed only modest effects of patient age on the CRC methylome after adjustment for confounders.

**Conclusions:** We identified a distinct methylation profile distinguishing MSS and MSI CRC, including *MLH1*-independent markers of MMRd, as well as novel differentially methylated loci within MSI subgroups. We further showed that anatomical location has a strong independent impact on the CRC methylome. Together, these findings refine the molecular characterisation of CRC and highlight potential epigenetic markers that could inform patient stratification and precision oncology.

## Introduction

Colorectal cancer (CRC) is the second leading cause of cancer-related mortality worldwide (Bray *et al*., 2024). In addition to genetic drivers, epigenetic dysregulation is a defining molecular feature of CRC that shapes tumour initiation and evolution (Hanahan, 2022; Heide *et al*., 2022). DNA methylation, the addition of a methyl group to cytosine-phosphate-guanine (CpG) dinucleotides, is a key epigenetic mechanism regulating gene expression and is frequently disrupted in cancer (Saghafinia *et al*., 2018).

Approximately 15–20% of CRCs display concordant hypermethylation at CpG islands against a background of global hypomethylation, a feature termed the CpG Island Methylator Phenotype (CIMP) (Toyota *et al*., 1999). CIMP-high (CIMP-H) tumours are associated with distinct clinicopathological features including *BRAF* mutation, older age and proximal colon location (Weisenberger *et al*., 2006). A common downstream consequence of CIMP is epigenetic silencing of *MLH1* through promoter hypermethylation, leading to mismatch repair deficiency (MMRd) and microsatellite instability (MSI) (Bateman, 2021). While CIMP-driven *MLH1* hypermethylation represents the most common mechanism of MMRd in sporadic CRC, a subset of tumours develop MMRd through biallelic somatic mutations in MMR genes in the absence of a global methylator phenotype.

Since CIMP, *MLH1* promoter hypermethylation and MMRd are highly correlated molecular features, most methylation changes observed in sporadic MMRd CRC are thought to reflect upstream CIMP-driven CpG island hypermethylation. However, it remains unclear whether *MLH1* silencing or MMRd are associated with additional, more specific methylation alterations independent of the broader methylator phenotype. Distinguishing methylation changes attributable to CIMP or *MLH1* silencing from those that accompany MMRd independently is therefore necessary to determine the extent to which MMRd contributes to epigenetic heterogeneity in CRC. A more refined molecular characterisation of methylation changes associated with CIMP, *MLH1* silencing and MMRd may improve understanding of inter-tumour variability in tumour biology and immunotherapy response, and advance precision oncology approaches in CRC (Ambrosini *et al*., 2025).

In this study, we performed comprehensive epigenome-wide profiling of DNA methylation in sporadic CRC. Specifically, we sought to (1) characterise methylation differences associated with MMRd after accounting for CIMP status and *MLH1* promoter hypermethylation, (2) isolate methylation changes associated with *MLH1* silencing independent of the global methylator phenotype, and (3) evaluate the influence of anatomical location and patient age on the CRC methylation landscape. Our findings provide new insights into the molecular heterogeneity of CRC and identify potential epigenetic markers with clinical relevance for patient stratification and therapeutic decision-making.

## Methods

### Ethics approval

Genomic and phenotypic data used in this study were obtained as part of the 100,000 Genomes Project (100kGP) study from the National Genomic Research Library (Genomics England), a research resource with approval to operate as a Research Tissue Bank by the HRA/Cambridge Central Research Ethics Committee (REC reference 20/EE/0035), with the principles of the Helsinki Declaration embedded into day-to-day operations. Data from the National Genomic Research Library can be accessed from within the secure Genomics England Research Environment, subject to institutional agreements. Normal tissue samples were collected in accordance with ethical guidelines approved by the South Central (Oxford B) Research Ethics Committee (REC reference 05/Q1605/66) and the HRA/Cambridge Central Research Ethics Committee (REC reference 20/EE/0035).

### Study cohort and clinical data

The study cohort comprised 272 primary fresh-frozen CRC tissue samples and 5 normal tissue controls. Whole-genome sequencing (WGS) and clinical data, including patient sex, age at sampling and tumour anatomical location were obtained through the Genomics England (GEL) Research Environment (RE) as part of the 100kGP main programme v18 release. Participants with Lynch syndrome were identified by the presence of a pathogenic germline variant in an MMR gene (*MLH1*, *MSH2*, *MSH6*, *PMS2*) using WGS data and were excluded from the study cohort. Tumour anatomical location information was derived from the av_tumour table within the RE and categorised as proximal (right-sided), including the caecum, ascending colon, hepatic flexure, and transverse colon; distal (left-sided), including the splenic flexure, descending colon, sigmoid colon; or rectal, including the rectosigmoid junction and rectum. For EWAS analysis, anatomical location was dichotomised into proximal and distal (including rectal). Microsatellite instability (MSI) status was determined using Detecting Microsatellite Instability by Next-Generation Sequencing (mSINGS) (Salipante et al., 2014), run with default settings on tumour WGS data aligned to GRCh38 within the RE. Samples with missing MSI status or anatomical location data were excluded, resulting in a final cohort of 259 CRC and 5 normal tissue samples. A flow diagram of the study design and sample inclusion criteria is shown in Figure 1.

**Fig. 1.**
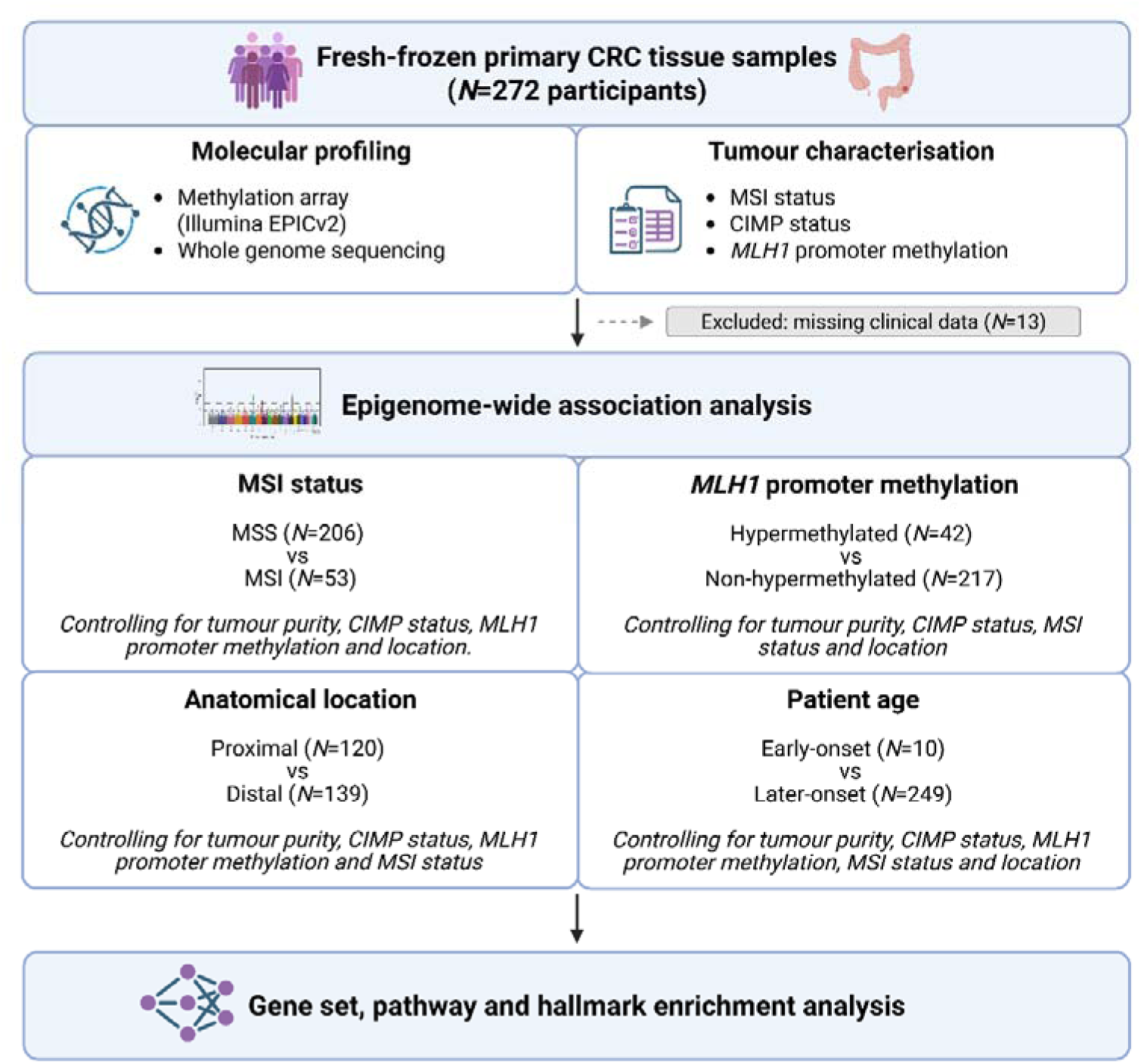
Study design. Flow diagram showing sample inclusion criteria and overview of the epigenome-wide association analyses performed. CRC, colorectal cancer; CIMP, CpG island methylator phenotype; MSS, microsatellite stable; MSI, microsatellite instability.

### DNA methylation data generation

DNA methylation data was generated from fresh-frozen tissue samples using the Illumina Infinium MethylationEPIC v2.0 BeadChip, which measures methylation at 937,691 CpG sites across the genome. For each sample, 500ng DNA was bisulphite-converted using the EZ DNA methylation-Gold kit (Zymo Research), according to the manufacturer’s protocol. To minimise batch effects, samples were randomised across bisulphite conversion plates, BeadChips and array positions.

### Data preprocessing and quality control

Data preprocessing, including sample-level and CpG site-level quality control was performed according to the following previously published pipeline (Hannon et al., 2024; https://github.com/ejh243/BrainFANS). Briefly, raw array IDAT files were imported into R using the bigmelon package (version 1.31.1) (Gorrie-Stone *et al*., 2019) and quality control checks were conducted using the wateRmelon package (version 2.11.4) (Pidsley *et al*., 2013). Poor quality samples were excluded using the following criteria; (i) median signal intensity of < 500, (ii) median bisulphite conversion efficiency < 80%, (iii) >1% probes with detection p > 0.05, (iv) samples with missing data for > 2% of probes, (v) normalised violence < 0.1 (wateRmelon qual() function, (vi) sample identity mismatches based on 65 SNP targeting probes. Poor quality probes were excluded using the following criteria; (i) detection P > 0.05 in more than 1% of samples, (ii) probes with common SNPs (minor allele frequency >5%) in the single base extension position based on those identified by Lehne et al. (Lehne *et al*., 2015), (iii) mis-mapped or mapping to non-specific regions.

Methylation levels at each CpG were defined as beta-values (β-values), which were calculated as the ratio of methylated signal intensity to the sum of the methylated and unmethylated signal intensity at each probe. To correct for technical variation between probes, and ensure comparability across samples, raw signal intensities were normalised using the Dasen function within the wateRmelon package.

### Tumour purity estimation

Tumour purity was estimated from the normalised β-values using InfiniumPurify (version 1.3.1) (Qin *et al*., 2018). Tumour cell fraction was inferred with the getPurity() function using informative differentially methylated CpG sites (iDMCs) derived from TCGA colon adenocarcinoma as the reference dataset by specifying tumour.type = “COAD”.

### Classification of *MLH1* promoter methylation

We selected four CpG probes, located within the *MLH1* promoter, based on the critical region defined by Deng et al (Deng *et al*., 2002). To classify promoter hypermethylation, we adapted the method described by Benhamida et al. for the Infinium MethylationEPIC v1.0 array (Benhamida *et al*., 2020). Three of the four probes (cg23658326, cg21490561, and cg00893636) used are present on both v1.0 and v2.0 arrays and were validated by Benhamida et al. The fourth probe, cg19132762, is specific to the v2.0 array and was included because of its proximity to the critical promoter region (22 bp upstream). Methylation thresholds for each probe were calculated as the mean β-value of normal tissue controls plus three standard deviations. Samples were classified as *MLH1-*hypermethylated if the β-value at each of the four probes exceeded the methylation threshold. Samples with β-values below the threshold for three or more probes were classified as non-*MLH1-*hypermethylated. Threshold values for each probe are provided in Supplementary Table 1.

### Classification of CIMP status

CIMP status was defined using the recursively-partitioned mixture model (RPMM) clustering method applied in previous CRC methylation studies (Muzny *et al*., 2012; Guinney *et al*., 2015; Ward *et al*., 2025). Briefly, RPMM clustering (RPMM R package, version 1.25) was performed on the 10% most variable autosomal probes (n = 84,886), resulting in CRCs being assigned into one of four methylation clusters, CIMP-High, CIMP-Low, Cluster 3 and Cluster 4, with Cluster 3 and Cluster 4 collectively described as “Non-CIMP”.

### Principal components analysis

Principal Component Analysis (PCA) was performed on the normalised β-values using the prcomp() function in R. To account for differences in DNA methylation across the genome, β-values for each CpG site (variable) were centred to a mean of zero and scaled to unit variance (Z-score transformation), prior to singular value decomposition. The proportion of variance explained by each principal component (PC) was evaluated using a scree plot (Supplementary Figure 1). Pearson correlation coefficients were computed between PCs and known clinical or technical covariates to identify potential confounders for inclusion in downstream regression models.

### Differential methylation analysis

Differentially methylated CpG sites were identified using the limma R package (version 3.64.1) (Ritchie *et al*., 2015), which fits a linear model for each CpG site to test the association between methylation and the variable of interest. The models were adjusted for potential confounders identified through PCA. Limma applies empirical Bayes moderation to the standard errors of the estimated log-fold changes, which enhances statistical power by borrowing information across all probes, yielding more stable and reliable variance estimates. Since β-values were used as input, the estimated effect sizes correspond to Δβ (difference in mean β-values) between groups. Significance was defined using a Bonferroni-adjusted *p*-value threshold of p<5.91x10⁻⁸ (to account for 846,923 individual probe-level tests).

### Genomic annotation of CpG sites

CpG sites were annotated to genes using a modified version of the Infinium MethylationEPICv2.0 manifest, re-annotated based on the GENCODE Human Release 47 (GENCODE v47) reference database (Mallabar-Rimmer *et al*., 2025). The genomic location of CpG sites relative to CpG islands was also classified as; CpG Islands (CGI), Shore (within 0–2 kb of a CpG island), Shelf (within 2-4 kb of a CpG island) and non-CGI-related sites.

### Gene set, pathway and hallmark enrichment analysis

To determine whether significantly differentially methylated CpG sites were enriched in specific biological pathways, functional enrichment analysis was performed using the gometh() function within the missMethyl R package (version 1.42.0) (Phipson, Maksimovic and Oshlack, 2016). This method accounts for probe-density bias due to differences in CpG representation across genes on the EPICv2 array. Enrichment was tested using the Gene Ontology (GO) and Kyoto Encyclopaedia of Genes and Genomes (KEGG) pathway databases. Functional profiling of Hallmark gene sets against the Molecular Signatures Database (MSigDB, version 7.1) was conducted using the gprofiler2 R package (version 0.2.3). Pathways were considered significantly enriched at a False Discovery Rate (FDR) < 0.05.

### Analysis of promoter methylation at CRC driver genes

To investigate methylation changes with potential functional impact on gene regulation in CRC, we performed a focused analysis on promoter regions of CRC driver genes. A curated list of 82 CRC driver genes was obtained from the Integrative OncoGenomics (IntOGen) database (version 2024.09.20) (Gundem *et al*., 2010), which identifies driver genes by aggregating and analysing somatic mutation data across multiple cancer types. Promoter-associated CpG sites were defined as those annotated as within 200 base pairs upstream of the transcriptional start site (TSS200), using GENCODEv47. Differential methylation analysis was performed on the subset of promoter-associated CpG sites in CRC driver genes using limma as described previously. A Bonferroni-adjusted *p*-value threshold of *p*<1.04×10⁻^4^ was applied to account for 479 probes.

### Data handling and statistical analysis

Methylation data preprocessing and quality control were performed using R (v4.2.1), functional enrichment analysis was conducted using R (v4.5), all other analyses were performed using R (v4.3.3). Summary statistics for cohort clinical variables were generated separately for MSS and MSI groups (Table 1). Categorical variables were reported as counts and percentages, and between-group differences were assessed using Fisher’s exact test. Continuous variables were summarised as mean ± standard deviation and compared using Welch’s *t*-test. For methylation analyses, Bonferroni correction was applied to adjust for multiple testing in the limma regression models. All other statistical tests were two-sided, and a *p*-value < 0.05 was considered statistically significant.

**Table 1.** Summary of the study cohort clinical and molecular characteristics. Counts <5 have been masked to preserve participant confidentiality. Comparisons between MSI and MSS groups were performed using Fisher’s exact tests for categorical variables and Welch’s two-sample *t*-test for continuous variables. CIMP, CpG island methylator phenotype; MSI, microsatellite unstable; MSS, microsatellite stable; SD, standard deviation.

| Characteristic | MSS | MSI | P value |
| --- | --- | --- | --- |
| **Total (N)** | 206 | 53 |  |
| **Age (mean ± SD)** | 69.5 ± 10.4 | 70.2 ± 11.7 | *p*=0.67 |
| **Sex** |  |  | *p*=3.59x10<sup>-5</sup> |
| Female | 67 (32.5%) | 34 (64.2%) |  |
| Male | 139 (67.5%) | 19 (35.8%) |  |
| **Anatomical location** |  |  | *p*=7.31x10<sup>-12</sup> |
| Proximal colon | 73 (35.4%) | 47 (88.7%) |  |
| Distal colon | 76 (36.9%) | ≤5 |  |
| Rectal | 57 (27.7%) | ≤5 |  |
| **MLH1 promoter status** |  |  | *p*=7.2x10<sup>-31</sup> |
| Hypermethylated | ≤5 | 39 (73.6%) |  |
| Non-hypermethylated | 203 (98.5%) | 14 (26.4%) |  |
| **CIMP status** |  |  | *p*=1.96x10<sup>-19</sup> |
| CIMP-High | 18 (8.7%) | 38 (71.7%) |  |
| CIMP-Low | 44 (21.4%) | ≤5 |  |
| Non-CIMP | 144 (69.9%) | 11 (20.8%) |  |
| **Stage** |  |  | *p*=0.436 |
| I | 28 (13.6%) | ≤5 |  |
| II | 74 (35.9%) | 25 (47.2%) |  |
| III | 57 (29.4%) | 16 (30.2%) |  |
| IV | 14 (6.8%) | ≤5 |  |
| Unknown | 33 (16%) | ≤5 |  |

## Results

### Cohort characteristics and principal determinants of methylation variance

We analysed epigenome-wide methylation and whole-genome sequencing data from 259 primary CRC cases, comprising 206 MSS and 53 MSI tumours (Figure 1). Clinical and molecular characteristics are summarised in Table 1. Consistent with previous studies, MSI CRC was more frequent in female patients (64.2% vs 32.5%, *p* = 3.59x10^-5^; Fisher’s exact test) and showed a strong association with anatomical location (*p* = 7.31x10^-12^; Fisher’s exact test), with 88.7% of MSI CRCs arising in the proximal colon compared with 35.4% of MSS CRCs (Figure 2B). As expected, MSI status was significantly associated with both *MLH1* promoter hypermethylation (73.6% of MSI vs <5% of MSS CRC, *p* = 7.2x10^-31^; Fisher’s exact test) and CIMP status (*p* = 1.96x10^-19^; Fisher’s exact test), with 71.7% of MSI CRC classified as CIMP-High compared to 8.7% of MSS CRC.

**Fig. 2.**
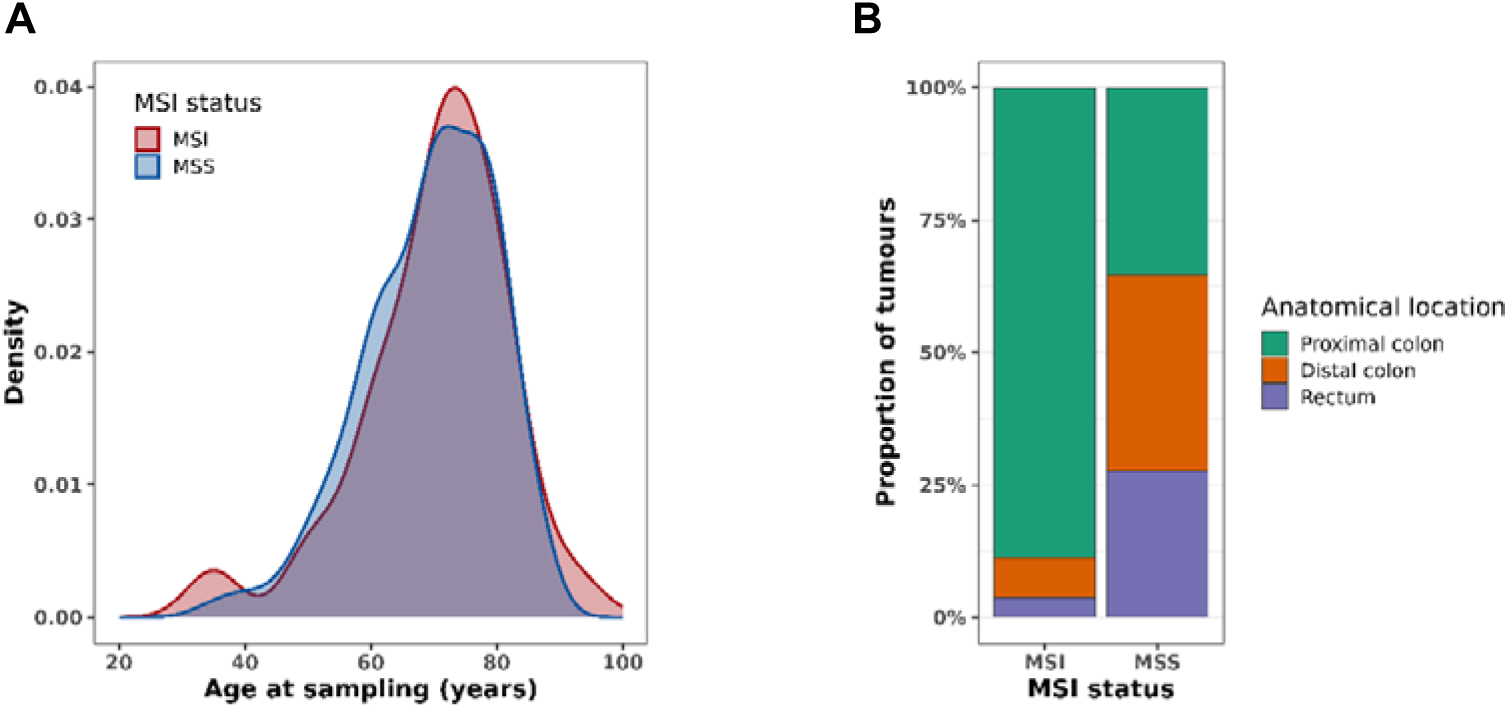
Clinical characteristics of colorectal cancer cohort stratified by microsatellite instability status (n=206 MSS; n = 53 MSI). (A) Density plot showing the age distribution at sampling for MSI and MSS samples. (B) Stacked bar chart showing relative proportions of MSI and MSS samples across anatomical locations of the colorectum. MSS, microsatellite stable; MSI, microsatellite instability.

We performed PCA on normalised β-values to identify technical and biological sources of methylation variance. Correlation of principal components (PCs) with clinical and molecular features revealed that MSI status, tumour purity, anatomical location, and *MLH1* promoter methylation were strongly associated with the first two PCs, each explaining >5 % of the total variance (Supplementary Figures 1-2). To account for potential confounding, these variables were included as covariates in downstream regression models when not the primary variable of interest. PCA score plots (PC1–PC2) coloured by key variables showed clean separation of MSI and MSS CRC along the first component, consistent with widespread methylation reprogramming associated with MMRd (Supplementary Figure 3).

### Epigenome-wide methylation differences between MSS and MSI CRC

An initial epigenome-wide association study (EWAS) comparing MSI (n=53) and MSS (n=206) CRC, without controlling for CIMP or *MLH1* promoter methylation status, identified 22,975 significantly differentially methylated positions (DMPs) (Bonferroni-corrected threshold *p* < 5.91x10⁻⁸, inflation λ = 4.11), confirming extensive global methylation differences between MSS and MSI CRCs (Supplementary Figure 4 and Supplementary Table 2). As anticipated, the signal was dominated by CpG sites at the *MLH1* promoter locus (Figure 3A). In addition to DMPs located in the *MLH1* promoter, we also observed other notable hypermethylated loci, including four consecutive significant DMPs annotated to *PRKCE* (median Δβ = 0.31, IQR: 0.29–0.33), highlighting additional loci that may be involved in the broader methylation changes characteristic of *MLH1*-hypermethylated MSI CRC.

**Fig. 3.**
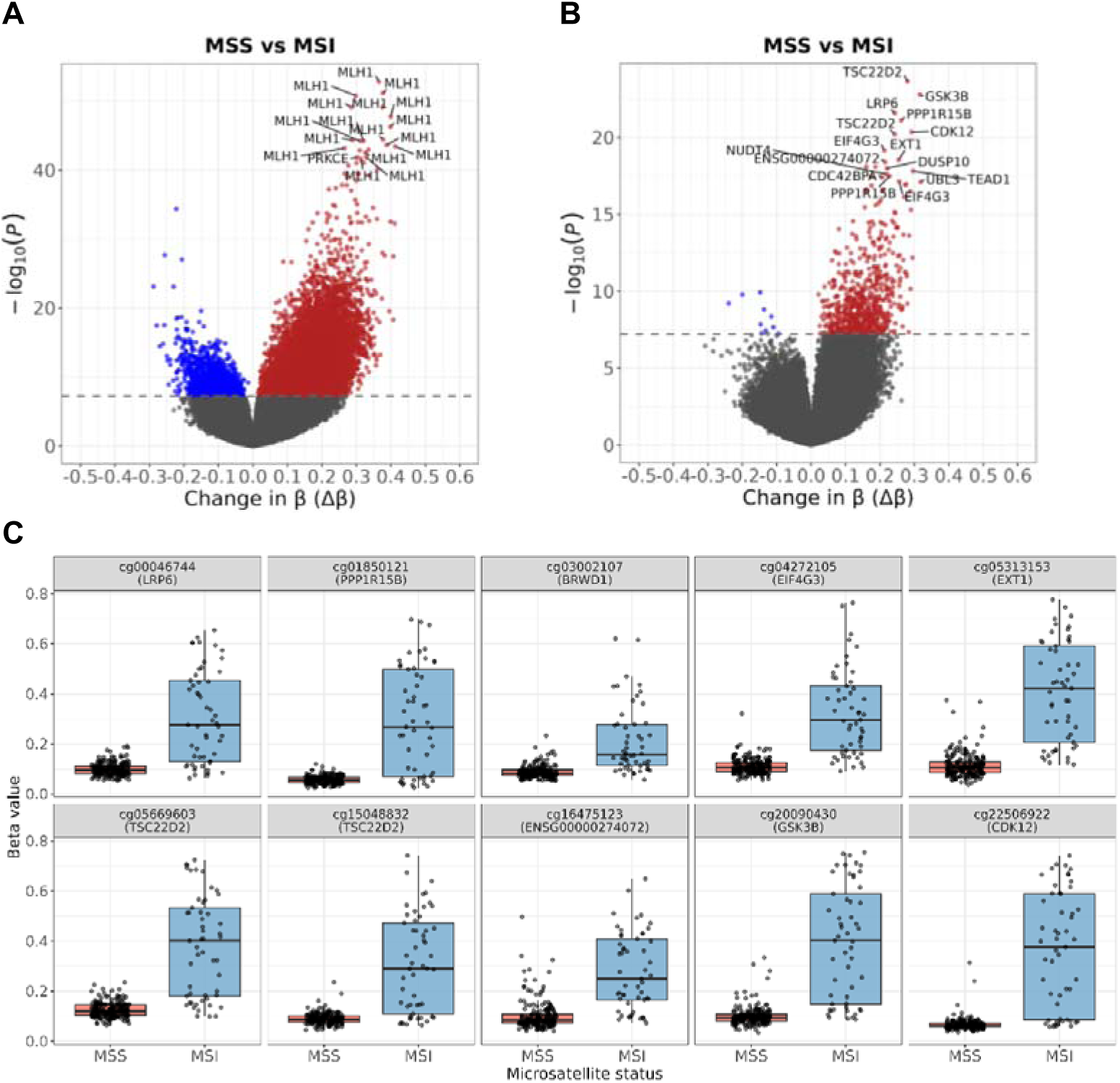
Epigenome-wide association study of DNA methylation in microsatellite stable (MSS; n=206) and microsatellite unstable (MSI; n=53) colorectal cancer. Volcano plots showing the association between methylation (difference in average beta, Δβ) and statistical significance (−log□ p-value) for (A) MSS vs MSI adjusted for tumour purity and anatomical location only, and (B) MSS vs MSI adjusted for tumour purity, anatomical location, *MLH1* promoter methylation and CIMP status. Each point represents a CpG site, top differentially methylated CpGs with Δβ > 0.15 are labelled with gene annotations. (C) Boxplots showing beta-values for the top 10 significantly differentially methylated CpG sites between MSS and MSI tumours, after adjustment for tumour purity, anatomical location, *MLH1* promoter methylation and CIMP status. These CpG sites represent the strongest epigenetic differences independent of the CIMP/*MLH1*-hypermethylated pathway and highlight potential methylation markers of mismatch repair deficiency. CIMP, CpG island methylator phenotype.

### Methylation differences associated with MMRd after accounting for CIMP and *MLH1* silencing

To disentangle the independent contribution of MMRd from effects of *MLH1*-hypermethylation or CIMP, we performed a subsequent analysis adjusting for both CIMP status (categorised as CIMP-H and CIMP-Low/negative) and *MLH1* promoter methylation in addition to tumour purity and anatomical location. We identified a substantially smaller set of 656 significant DMPs (*p* < 5.91×10⁻⁸, Supplementary Table 3) and a marked reduction in test statistic inflation (λ = 1.37 vs 4.11), representing methylation differences that are independent of the broader methylator phenotype (Supplementary Figure 5).

The majority (96.9%) of significant DMPs were hypermethylated in MSI relative to MSS CRC (Figure 3B). Of the most significantly associated DMPs, a prominent subset mapped to genes involved in WNT signalling including *LRP6*, a WNT co-receptor; *GSK3β*, a regulator of β-catenin stability; and *EXT1*, a heparan-sulphate polymerase involved in ligand presentation (Figure 3C). Further notable hypermethylated DMPs annotated to genes involved in TGF-β signalling (including *TSC22D2*, *TEAD1* and *SMAD3*) and genes related to transcriptional or RNA regulatory processes (*CDK12*, *EIF4G3* and *PPP1R15B*). To determine whether these MMRd-associated loci converge on specific biological processes, we performed pathway enrichment analysis of significant DMPs using Gene Ontology (GO), Kyoto Encyclopaedia of Genes and Genomes (KEGG), and Hallmark databases. No pathways or gene ontology terms reached statistical significance after multiple-testing correction (FDR < 0.05, Supplementary Table 4), suggesting that methylation changes associated with MMRd are not clustered within canonical pathways.

Together, these findings demonstrate that, after accounting for the effects of the CIMP and *MLH1* promoter hypermethylation, only a limited residual set of methylation differences distinguishes MSS and MSI subtypes. The marked attenuation of both the number of significant loci and test statistic inflation after adjustment indicates that the majority of methylation differences between MSI and MSS CRC reflect upstream CIMP-related processes, with only a modest residual signal associated with MSI status.

### Methylation differences associated with *MLH1* promoter hypermethylation after accounting for CIMP

*MLH1* promoter hypermethylation, arising as a downstream consequence of CIMP, is the primary epigenetic mechanism leading to sporadic MMRd in CRC. However, *MLH1* promoter hypermethylation may also function as a distinct epigenetic event associated with methylation changes in addition to those of the broader methylator phenotype. We therefore sought to characterise methylation patterns accompanying *MLH1* promoter hypermethylation by comparing *MLH1*-hypermethylated and non-*MLH1*-hypermethylated tumours, adjusting for tumour purity, anatomical location, and CIMP status. This analysis identified 3,508 significant DMPs (*p* < 5.91 x 10⁻⁸, inflation λ = 1.33, Supplementary Table 5), reflecting widespread background hypermethylation characteristic of CIMP (Supplementary Figure 6). As expected, the most significant associations mapped to the *MLH1* promoter region. Several additional loci showed concurrent hypermethylation in *MLH1*-hypermethylated tumours, including *B3GALT1* and *SCNNB1* (*p* < 5.91 x 10⁻⁸). Gene set and pathway enrichment analysis of *MLH1*-associated differentially methylated positions (DMPs) highlighted genes involved in developmental and neurogenic processes, as well as ion channel activity (FDR < 0.05, Supplementary Figure 7A and Supplementary Table 6). Enriched pathways included neuroactive ligand–receptor interactions, neuroactive ligand signalling and calcium signalling (Supplementary Figure 7B), suggesting that *MLH1*-associated methylation changes may influence cell signalling beyond canonical DNA repair pathways.

### Co-methylation of *MLH1* with *TXNRD1* in MSI CRC

To distinguish methylation changes driven by *MLH1* silencing from those arising through alternative MMRd mechanisms, we directly compared *MLH1*-hypermethylated (n=39) and non-*MLH1*-hypermethylated (n=14) MSI tumours, adjusting for tumour purity, anatomical location and CIMP status (Figure 4A). We identified 81 significant DMPs mapping to 29 unique genes (*p* < 5.91×10⁻⁸; λ = 0.97, Supplementary Table 7). In addition to the *MLH1* locus, top DMPs were annotated to *TXNRD1,* a thioredoxin reductase enzyme involved in the regulation of cellular oxidative stress, *CYP1B1*, and *RERG*. Five hypermethylated CpG sites were located within the *TXNRD1* promoter, and *MLH1* promoter methylation was strongly correlated with *TXNRD1* promoter methylation (Pearson’s *r* = 0.98, *p* < 8.1×10⁻³⁷, Figures 4B–C), consistent with co-occurring promoter hypermethylation in *MLH1*-hypermethylated MSI. Although significant DMPs were distributed across multiple genes, enrichment analyses did not yield any significantly enriched GO, KEGG or Hallmark terms (FDR > 0.05; Supplementary Table 8).

**Fig. 4.**
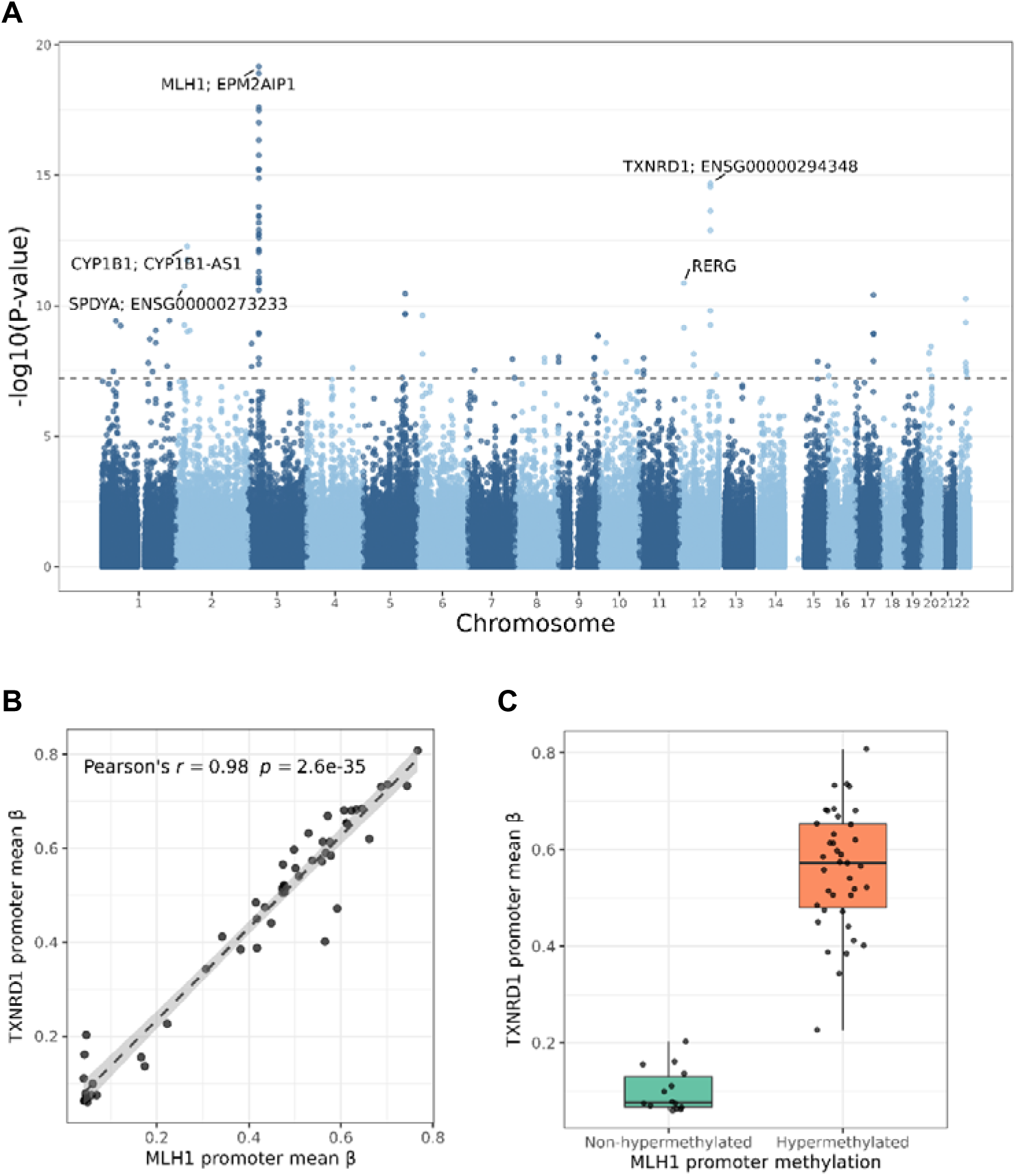
Epigenome-wide association of microsatellite unstable (MSI) colorectal cancer (CRC) with (n =39) and without (n =14) *MLH1* promoter hypermethylation, adjusting for tumour purity, anatomical location and CIMP status. (A) Manhattan plot showing −log□□ p-values for all CpG sites, Bonferroni-corrected epigenome-wide significance (p< 5.91 x 10^−^□) is indicated by the grey dashed line. (B, C) Scatter plots showing the relationship between mean promoter methylation (CpG sites annotated to TSS200) in *MLH1* and *TXNRD1*. Dashed lines represent linear regression fits, with correlation (r) calculated using Pearson’s correlation. (C) *TXNRD1* promoter methylation by *MLH1* promoter hypermethylation status. Boxplots showing beta values for TSS200 CpGs in *TXNRD1* across *MLH1*-hypermethylated versus non-hypermethylated MSI CRC.

### Impact of anatomical location and patient age on DNA methylation profiles

PCA showed that anatomical location was a major source of variation in global CRC methylation profiles. We next assessed methylation differences by anatomical location, comparing proximal (n=120), and distal (n=139) CRCs, adjusting for tumour purity, CIMP status, *MLH1* promoter methylation and MSI status. We identified 1,428 significant DMPs (*p* < 5.91×10⁻⁸, inflation λ = 2.14, Supplementary Figure 8A), annotated to 727 unique genes, indicating marked differences in methylation with anatomical location (Figure 4A, Supplementary Table 9). The majority (89.8%) of significant DMPs were hypermethylated in proximal CRC. Among the most significant DMPs were CpG sites annotated to *CDX2*, a homeobox intestinal transcription factor critical for intestinal differentiation; *CTNND2*, a known oncogenic driver; *FMN2*, a putative tumour suppressor; and *PRAC1*, *PDE10A* and *ELAVL2* (Figure 5B). Proximal CRCs showed coordinated hypomethylation of five consecutive CpG sites in the *CDX2* promoter region (TSS1500), and twelve adjacent CpG sites within the promoter of *PDE10A*. In contrast, *FMN2*, *PRAC1/2* and *ELAVL2* showed consistent hypermethylation across multiple promoter-associated CpG sites (Figure 5C). These results demonstrate that anatomical location is a major determinant of CRC methylation patterns, independent of MSI or CIMP status and *MLH1* promoter methylation.

**Fig. 5.**
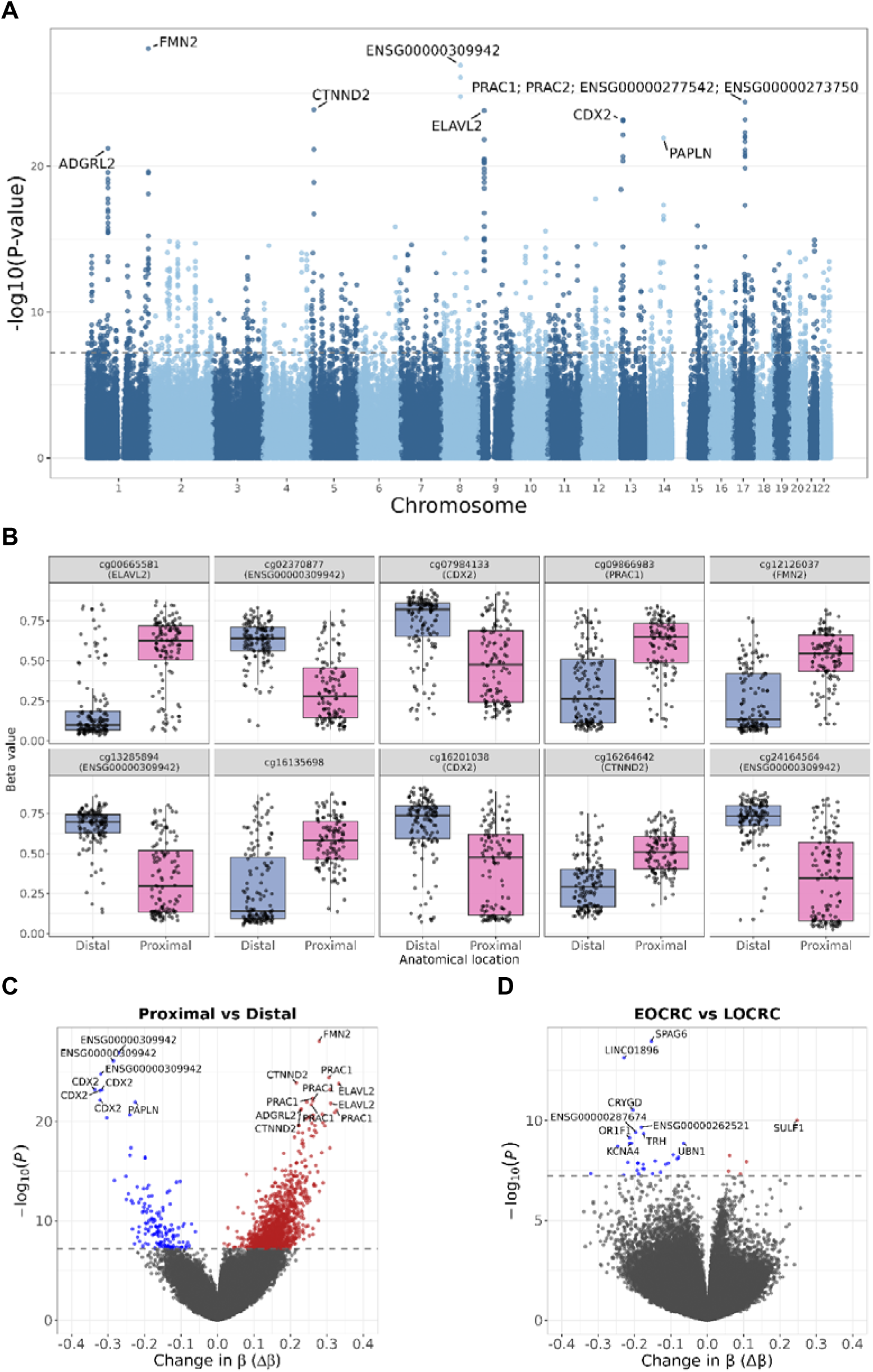
Epigenome-wide association analysis of anatomical location and patient age in colorectal cancer CRC. (A) Manhattan plot showing −log□ p-values for differentially methylated CpG sites between proximal (n =120) and distal (n =139) CRC, adjusted for tumour purity, MSI status, CIMP status, and *MLH1* promoter hypermethylation. The grey dashed line indicates the Bonferroni-corrected epigenome-wide significance threshold (p < 5.91 × 10□). (B) Boxplots showing the distribution of β-values for the top 10 significantly differentially methylated CpG sites between proximal and distal CRC. (C) Volcano plot showing methylation differences (Δβ) against statistical significance (−log□ p-value) for proximal vs distal CRC. (D) Volcano plot showing methylation differences for early-onset CRC (n=10) vs later-onset CRC (n = 249). Genes annotated to the most significant differentially methylated CpGs are labelled. CRC, colorectal cancer; MSI, microsatellite instability; CIMP, CpG island methylator phenotype.

Gene ontology analysis of significant DMPs revealed strong enrichment for developmental and morphogenetic processes, consistent with the distinct embryonic origins of the proximal (midgut) and distal (hindgut) colon (Supplementary Figure 9 and Supplementary Table 10). Top enriched terms included multicellular organism development, system development, and anatomical structure morphogenesis (FDR < 0.05). Enriched molecular function categories highlighted regulators of chromatin and transcription, including DNA-binding transcription factor activity and RNA polymerase II cis-regulatory region binding, indicating preferential involvement of promoter- and enhancer-associated regulatory regions.

We next evaluated the contribution of patient age to methylation variation. Using age as a continuous variable and adjusting for MSI status, tumour purity, anatomical location, CIMP status, and *MLH1* promoter methylation, we did not identify any significant DMPs at the Bonferroni-adjusted threshold (p < 5.91×10⁻⁸, inflation λ = 0.92, Supplementary Figure 10 and Supplementary Table 11). In contrast, when modelling age as a binary variable comparing early-onset CRC (EOCRC, <50 years, n=10) and later-onset CRC (LOCRC, ≥50 years, n=249), we identified 34 significant DMPs (Supplementary Table 12). The most significant loci mapped to genes including *SPAG6*, *LINC01896*, *SULF1*, and *CRYGD* (Figure 5D).

### Analysis of promoter methylation at CRC driver genes

To investigate whether age or anatomical location influences the epigenetic regulation of known CRC driver genes, we performed a targeted analysis of 82 driver genes curated from the Integrative OncoGenomics (IntOGen) database, specifically focusing on CpG sites within the promoter regions (TSS200). Restricting differential methylation analysis to these regions, while controlling for tumour purity, CIMP status, *MLH1* promoter methylation and MSI, revealed 4 DMPs significantly associated with anatomical location after Bonferroni correction (*p*<1.04×10⁻^4^). Of the significantly associated CpG sites, 3 were located within the promoter region of *GRIN2A* and 1 within the promoter region of *ERBB2*, all of which were hypermethylated in proximal compared distal tumours (Supplementary Figure 11A, Supplementary Table 13). In contrast, no CpG sites within these driver gene promoters were significantly associated with patient age (Supplementary Figure 11B, Supplementary Table 14). These results suggest that specific epigenetic divergence in *GRIN2A* and *ERBB2* may reflect distinct oncogenic pathways between proximal and distal CRC, whereas patient age has limited impact on driver gene promoter methylation.

## Discussion

In this study, we integrate genomic and methylation data to provide a comprehensive characterisation of the epigenetic variation across molecular and clinical subtypes of CRC. We identify both established and novel methylation features associated with MMRd, anatomical location, and patient age. By accounting for major sources of methylation variation, including tumour purity, *MLH1* promoter methylation, CIMP status and site-specific effects, we distinguish distinct methylation changes specifically associated with MMRd, from those accompanying *MLH1* silencing or CIMP. Together, these findings refine the molecular characterisation of CRC subtypes and improve our understanding of the epigenetic mechanisms that shape CRC evolution.

Consistent with its established role as the primary driver of sporadic MMRd, an initial comparison of MSI and MSS CRC revealed extensive methylation differences dominated by hypermethylation at the *MLH1* promoter and widespread CIMP-associated changes. In addition to *MLH1* hypermethylation, we also observed significant hypermethylation at other genes, including *PRKCE*, a serine-threonine kinase reported to stabilise TET1 via protein phosphorylation (Chen *et al*., 2023). Although the high genomic inflation observed in this analysis warrants cautious interpretation, given the role of TET1 in active demethylation and its association with CIMP (Tricarico *et al*., 2023), *PRKCE* hypermethylation may represent a CIMP-related feature contributing to altered TET1 regulation.

After controlling for *MLH1* promoter methylation and CIMP status we identified a distinct set of DMPs associated with MMRd. The most significantly associated loci mapped to genes involved in WNT signalling, including *GSK3β*, *LRP6* and *EXT1* as well as regulators of transcription control such as *CDK12*, *TSC22D2* and *EIF4G3*. *GSK3β* promotes β-catenin degradation in the cytoplasmic destruction complex, *LRP6* functions as a cell surface co-receptor initiating ligand-dependent activation and *EXT1*, a heparan-sulphate polymerase required for WNT and FGF ligand presentation (Yamamoto *et al*., 2013; Nusse and Clevers, 2017). While aberrant WNT activation in CRC is typically driven by *APC* mutations, these findings suggest that altered epigenetic regulation of WNT signalling components may also contribute to pathway dysregulation in MMRd, although functional validation is required to establish whether these methylation differences directly modulate WNT activity.

Additionally, *CDK12* (cyclin-dependent kinase 12) regulates transcription of DNA damage repair genes (Krajewska *et al*., 2019). Loss or reduced expression of *CDK12* through promoter hypermethylation may contribute to genomic instability in non-*MLH1*-hypermethylated MSI CRC. Interestingly, hypermethylation at *TSC22D2*, encoding Transforming Growth Factor β- Stimulated Clone 22 Domain Family Member 2, has recently been reported as a robust biomarker for MSI detection and is positively associated with features predictive of immunotherapy response, including tumour mutational burden, neoantigen load, and expression of immune checkpoint molecules (Cao *et al*., 2023). Our independent identification of this locus provides further support for *TSC22D2* methylation as a clinically relevant epigenetic marker of MMRd.

Direct comparison of *MLH1*-hypermethylated and non-*MLH1*-hypermethylated MSI CRC revealed a small set of co-methylated loci, most notably including *TXNRD1*. The strong correlation between *TXNRD1* and *MLH1* promoter hypermethylation suggests coordinated epigenetic regulation in MSI CRC. *TXNRD1* encodes a thioredoxin reductase enzyme involved in mitigating the accumulation of reactive oxygen species (ROS) (Harris *et al*., 2015), suggesting a potential link between altered methylation patterns and oxidative stress pathways in MSI CRC. Since our analysis controlled for CIMP status, hypermethylation at *TXNRD1* and other significant loci likely reflects MSI-specific epigenetic alterations, rather than global CIMP- related hypermethylation. The consistent co-occurrence of *TXNRD1* and *MLH1* promoter hypermethylation therefore raises the possibility that impaired oxidative stress responses could represent a recurrent feature of *MLH1*-hypermethylation MSI CRC, and highlights *TXNRD1* as a potentially functionally relevant gene warranting further investigation.

Beyond differences between CRC molecular subtypes, we found a strong independent impact of anatomical location on the CRC methylome, identifying 1,428 significant DMPs between proximal and distal CRCs, after adjusting for MSI status, CIMP status, *MLH1*-hypermethylation, and tumour purity. Among the most significant loci, were CpG sites annotated to *CDX2, ELAVL2, PRAC1/2* and *PDE10A*. CDX2 is an intestinal transcription factor and tumour suppressor that maintains colonic epithelial differentiation and limits proliferation, partly through repression of Wnt/β-catenin signalling (Yu et al., 2019). We observed hypomethylation of CpG sites annotated to *CDX2* in proximal compared with distal CRC, consistent with previous findings in adenomas from the left and right colon (Koestler *et al*., 2014). While loss of expression of CDX2 has previously been associated with molecular features of the serrated pathway, MMRd, CIMP-H and poor prognosis (Dalerba *et al*., 2016; Graule *et al*., 2018; Yang *et al*., 2024), our results suggest that there are significant differences in methylation at this locus associated with anatomical origin, independent from the effects of CIMP. Conversely, we identified hypermethylation at multiple CpG sites at *PRAC1/2* and *ELAVL2* in proximal CRC. The hypermethylation of *PRAC1/2* suggests that DNA methylation may contribute to the reduced PRAC1 expression observed in proximal CRC in previous studies (Hu *et al*., 2018; Jiang *et al*., 2020; Kolisnik *et al*., 2023). Similarly, hypermethylation of the RNA-binding protein *ELAVL2*, is consistent with previously identified transcriptomic differences between left- and right-sided CRC (Du *et al*., 2021), suggesting location-specific silencing of regulatory genes in the proximal colon. Furthermore, the differential methylation of *PDE10A* is also of interest, as this gene is a known proto-oncogene in CRC and a potential therapeutic target.

In contrast to the strong influence of anatomical location, we observed only a modest impact of patient age at diagnosis on DNA methylation patterns in our cohort. When age was modelled as a continuous variable, no CpG sites reached epigenome-wide significance after multiple testing correction, suggesting that age-related methylation effects are not a dominant feature of the CRC methylome. Although age has been associated with both methylation changes in normal tissues (Teschendorff and Horvath, 2025), and with *MLH1* hypermethylation in CIMP-H CRC (Levine et al., 2016), our PCA and EWAS results suggest that age-related methylation changes may be less pronounced in tumour tissue relative to other tumour-associated epigenetic alterations. When age was instead modelled as a binary variable to comparing EOCRC (age < 50 years) and LOCRC (age ≥ 50 years), we identified 34 significant DMPs, including loci annotated to *SPAG6*, *LINC01896*, *SULF1*, and *CRYGD*. *SULF1* has recently been shown to be highly expressed in cancer-associated fibroblasts (Wang et. al 2024). These findings potentially reveal methylation differences reflecting distinct biological processes between EOCRC and LOCRC. However, the small size of the EOCRC group limits statistical power and independent validation in larger cohorts is required.

Through targeted analysis of promoter methylation in established CRC driver genes, we identified hypermethylation of three loci within the *GRIN2A* promoter in proximal CRC. Since our analysis controlled for CIMP status, *MLH1* promoter methylation, and MSI status, these results indicate that *GRIN2A* hypermethylation may be independent of the global CpG hypermethylation typically observed with CIMP. *GRIN2A* is a tumour suppressor, and its dysregulation has been previously linked to increased tumour recurrence (Metzger *et al*., 2025). While further functional validation would be required to confirm the impact on gene expression, these findings suggest that epigenetic silencing of *GRIN2A* may contribute to the distinct clinical trajectories observed in proximal CRC. In contrast, the absence of significant associations with patient age suggests that promoter methylation of these 82 CRC driver genes is not significantly influenced by chronological age in this cohort.

We acknowledge several limitations of this study. Firstly, the use of bulk tumour tissue precluded resolution of cell-type-specific methylation patterns. Although we adjusted regression models for tumour purity, this approach does not capture variations in cellular composition, such as differences in immune infiltration, which are known to vary between MSS and MSI subtypes and represent a common source of residual confounding in EWAS analyses (Zheng *et al*., 2017). As a result, some of the observed methylation signals associated with MMRd may partially reflect differences in immune or stromal cell proportions rather than tumour-intrinsic changes. Single-cell methylation profiling would offer higher-resolution insights into the cell-type specificity of the methylation changes we observed. Secondly, absence of matched normal tissue, limited interpretation of the tumour-specificity of methylation differences. Additionally, we did not account for prior treatments (e.g., chemotherapy, radiotherapy) and other exposures like smoking history, which are known to influence DNA methylation, and could introduce residual confounding (Yuan *et al*., 2025). Finally, while we have identified differential methylation signals, the functional impact of these differences on gene expression was not examined. Future studies integrating matched normal tissue, gene expression data (e.g., RNA-seq), and independent validation cohorts will be important to confirm the clinical relevance of the methylation changes identified.

## Conclusions

Here we define a distinct DNA methylation pattern specifically associated with MMRd in CRC, independent of *MLH1* promoter hypermethylation and broader CIMP-related effects. The most significant loci mapped to genes including *LRP6*, *GSK3β*, *CDK12*, and *TSC22D2*, implicating dysregulation of WNT signalling and transcriptional control as recurrent features of MMRd CRC. We further identify co-occurrence of promoter hypermethylation of *MLH1* and *TXNRD1* in MSI CRC, suggesting a novel link between MMRd and cellular oxidative stress regulation.

In addition to MMR status, we demonstrate that anatomical location is a major independent determinant of the CRC methylome, with proximal and distal CRC displaying distinct methylation profiles across developmental and regulatory genes including *CDX2*, *PRAC1/2*, and *PDE10A*. In contrast, we find only modest effects of patient age at diagnosis on methylation variation after adjustment for molecular and clinical covariates. Together, these findings refine the epigenetic classification of CRC, highlight novel candidate markers for molecular stratification, and provide a foundation for functional studies linking methylation patterns to CRC biology and therapeutic response.

## Acknowledgements

We gratefully acknowledge the participants of the National Genomic Research Library (NGRL), whose contributions made this research possible. Secure access to the NGRL under Research Registry Project ID; 1405: “The Epigenetic Landscape of Colorectal Cancer” was provided by Genomics England, which delivers the NGRL in partnership with NHS England, and is wholly owned by the UK Department of Health and Social Care. The NGRL contains participants’ health data collected by the NHS as part of their care, along with samples and data from their participation in research, for which fully informed consent has been obtained. This includes genomic and clinical data provided through the NHS Genomic Medicine Service, as well as data obtained through research studies, including the 100,000 Genomes Project and the Generation Study, both of which are delivered in partnership with the NHS, and from other research cohorts involving external collaborators. We are grateful to Dr Juan Fernández-Tajes for providing mSINGS data to derive the microsatellite instability status of samples and to Amy McCorry for assisting with the sample processing. This work was supported by the National Institute for Health and Care Research Exeter Biomedical Research Centre. The views expressed are those of the author(s) and not necessarily those of the NIHR or the Department of Health and Social Care.

## Data availability

Data from the National Genomic Research Library (NGRL) used in this research are available within the secure Genomics England Research Environment. Access to NGRL data is restricted to adhere to consent requirements and protect participant privacy. Data used in this research include: Clinical and genomic data from the 100kGP Main Programme Data Release v18.

Clinical data including patient age, sex and tumour anatomical location was extracted using the cancer_analysis and av_tumour tables. The full list of Genomics England participant IDs together with clinical information and scripts can be found at ‘re_gecip/cancer_colorectal/rward/Ward_et_al.2026_CRC_methylation’.

Access to NGRL data is provided to approved researchers who are members of the Genomics England Research Network, subject to institutional access agreements and research project approval under participant-led governance. For more information on data access, visit: https://www.genomicsengland.co.uk/research.

## Consent for publication

All participants in this study have provided consent for their data to be part of the National Genomics Research Library and to the publication of research findings. For all cases, written informed consent for research use of clinical and genetic data was obtained from patients, their parents, or legal guardians in the case of those with intellectual disability.

## Author contributions

Conceptualisation; R.W., A.P.W. Methodology; R.W., B.M.R, A.P.W. Software; R.W., M.E., B.M.R, Q.H., J.C.W., S.T., C.W., A.P.W. Validation; R.W., M.E., B.M.R, Q.H., J.C.W., S.T., C.W. Formal Analysis; R.W., M.E., Q.H., J.C.W. Investigation; J.B., K.S., S.J.W. Resources; I.T., A.P.W. Data Curation; R.W., M.E., Q.H., J.C.W., S.T., C.W. Writing – Original Draft; R.W. Writing – Review & Editing; R.W., M.E., B.M.R, K.S., Q.H., J.C.W., E.D., H.D.G., A.P.W. Visualisation; R.W., B.M.R, H.D.G., A.P.W. Supervision; E.D., H.D.G., I.T., A.P.W. Project Administration; M.E., I.T., A.P.W. Funding Acquisition; H.D.G., I.T., A.P.W.

