## Supplementary Figure for "DNA methylation signatures of mismatch repair-deficient colorectal cancer"

### Supplementary Figures

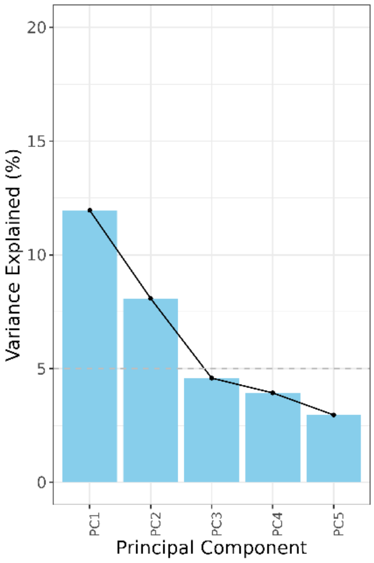


**Supplementary Fig. 1**. Scree plot showing the percentage of variance explained by the first 5 principal components (PCs), based on DNA methylation data from the full colorectal cancer cohort (n = 259).

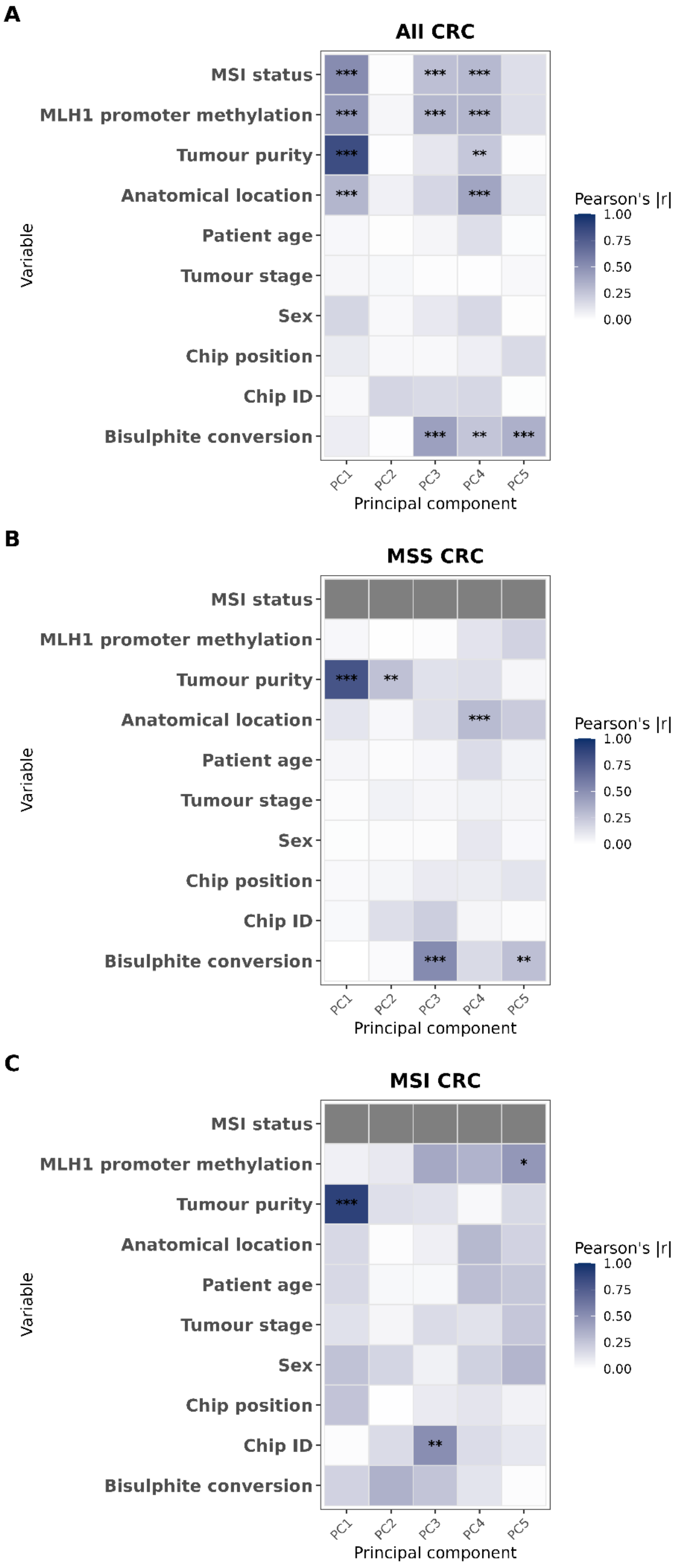
A
All CRC
B
MSS CRC
C
MSI CRC
MSI status
MLH1 promoter methylation
Tumour purity
Anatomical location
Patient age
Tumour stage
Sex
Chip position
Chip ID
Bisulphite conversion
Variable
PC1
PC2
PC3
PC4
PC5
Principal component
Pearson's |r|
1.00
0.75
0.50
0.25
0.00

**Supplementary Fig. 2**. Heatmap showing the absolute strength of correlation between principal components (PCs) derived from methylation beta values and clinical and technical variables in (A) all CRC samples (n=259), (B) microsatellite stable (MSS) CRC (n =206) and microsatellite unstable (MSI) CRC (n = 53). Each tile represents the absolute Pearson correlation coefficient (|r|) between a given PC and each variable. The tile colour indicates the strength of correlation (light blue = weaker, dark blue = stronger). Asterisks denote statistical significance after Bonferroni correction: ***p < 0.001, **p < 0.01, *p < 0.05.

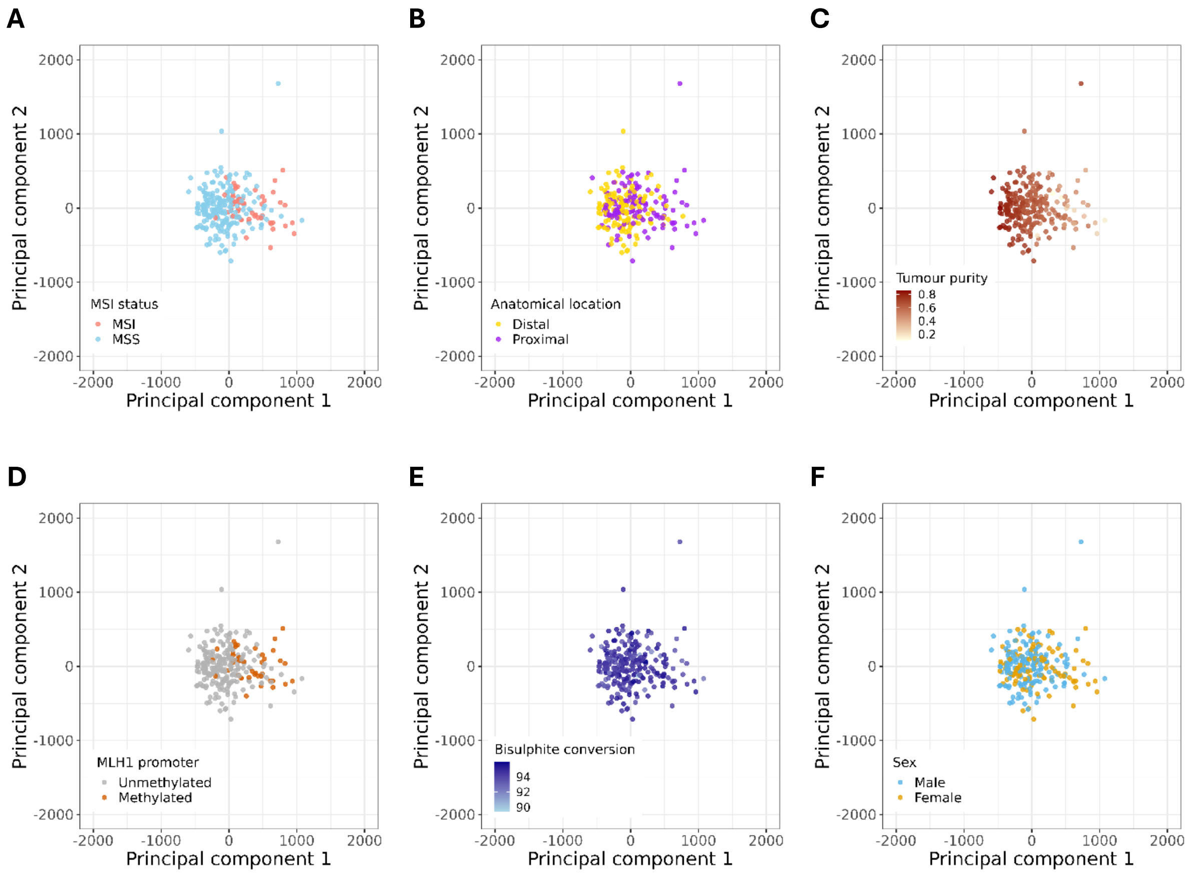


**Supplementary Fig. 3**. Principal component analysis (PCA) score plots of methylation β-values for all colorectal cancer samples (n=259). PC1 and PC2 coordinates are shown, with individual samples coloured by key clinical or technical variables. A) MSI status, B) anatomical location, C) tumour purity, D) *MLH1* promoter methylation status, E) bisulphite conversion efficiency and F) patient sex.

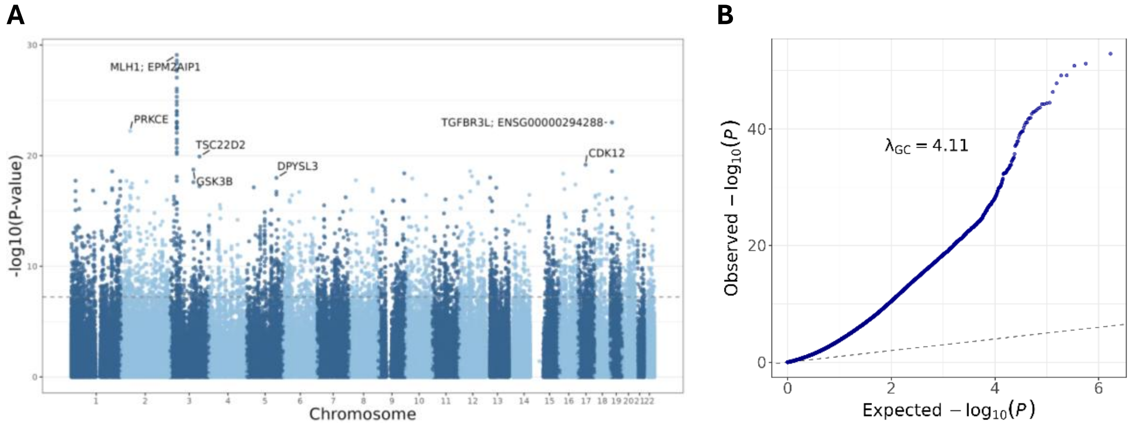


**Supplementary Fig. 4.** Quantile–quantile (QQ) and Manhattan plots from the epigenome-wide association between microsatellite stable (n=206) and microsatellite unstable (n=53) tumours adjusted for tumour purity and anatomical location (A) Manhattan plot showing $-\log_{10}$ p-values for all CpG sites across the genome. The grey dashed line indicates the Bonferroni-corrected epigenome-wide significance threshold ($p < 5.9 \times 10^{-8}$). Genes annotated to the most significantly differentially methylated CpG sites are labelled. (B) QQ plot showing observed versus expected $-\log_{10}$ p-values.

A

B

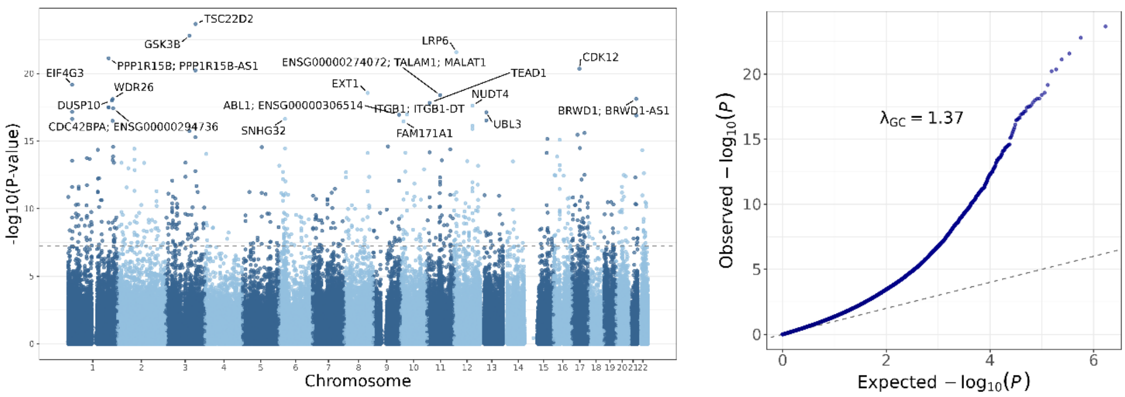


**Supplementary Fig. 5.** Manhattan and quantile-quantile (QQ) plots from the epigenome-wide association between microsatellite stable (n=206) and microsatellite unstable (n=53) tumours adjusted for tumour purity, anatomical location, CIMP status and *MLH1*-hypermethylation (A) Manhattan plot showing $-\log_{10}$ p-values for all CpG sites across the genome. The grey dashed line indicates the Bonferroni-corrected epigenome-wide significance threshold ($p < 5.9 \times 10^{-8}$). Genes annotated to the most significantly differentially methylated CpG sites are labelled. (B) QQ plot showing observed versus expected $-\log_{10}$ p-values.

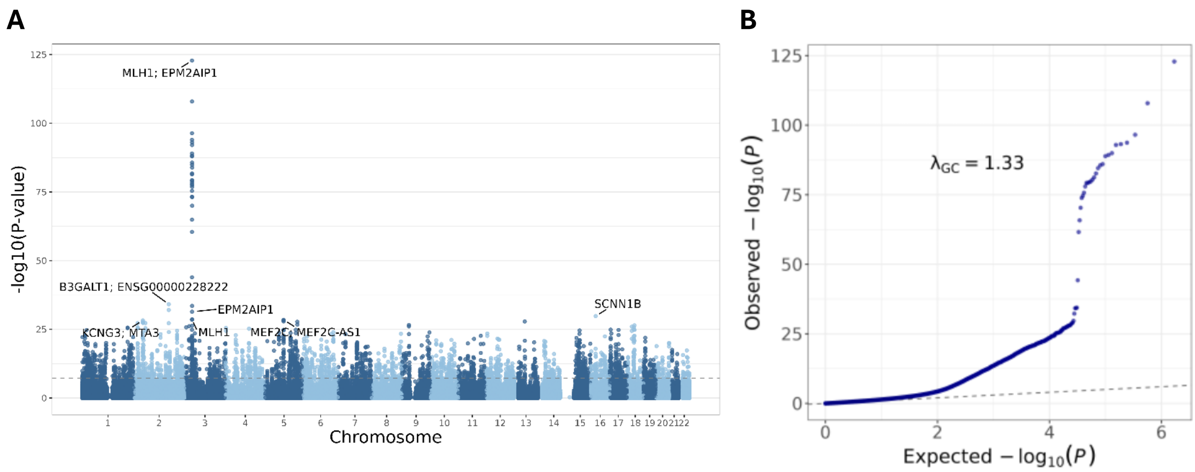


**Supplementary Fig. 6.** Manhattan and quantile–quantile (QQ) plots from the epigenome-wide association between *MLH1*-hypermethylated (n=42) and non-*MLH1*-hypermethylated (n=217) colorectal cancer samples, adjusting for tumour purity, anatomical location and CIMP status (A) Manhattan plot showing $-\log_{10}$ p-values for all CpG sites across the genome. The grey dashed line indicates the Bonferroni-corrected epigenome-wide significance threshold ($p < 5.9 \times 10^{-8}$). Genes annotated to the most significantly differentially methylated CpG sites are labelled. (B) QQ plot showing observed versus expected $-\log_{10}$ p-values.

**A**

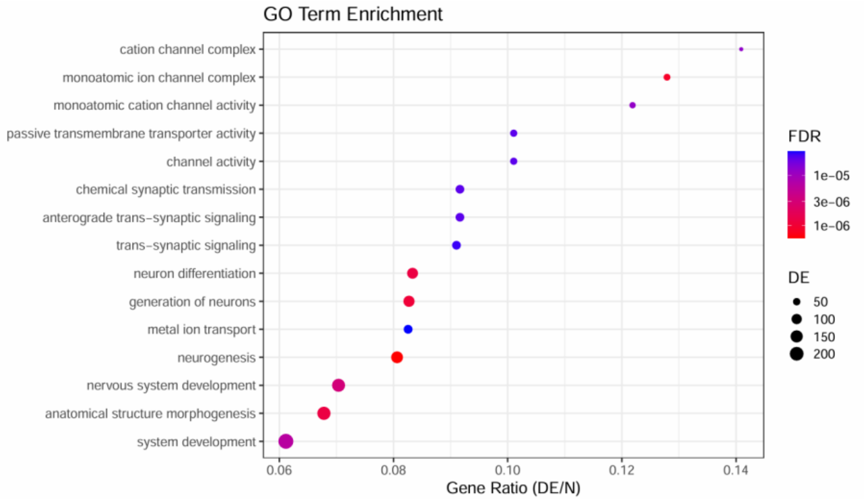


**B**

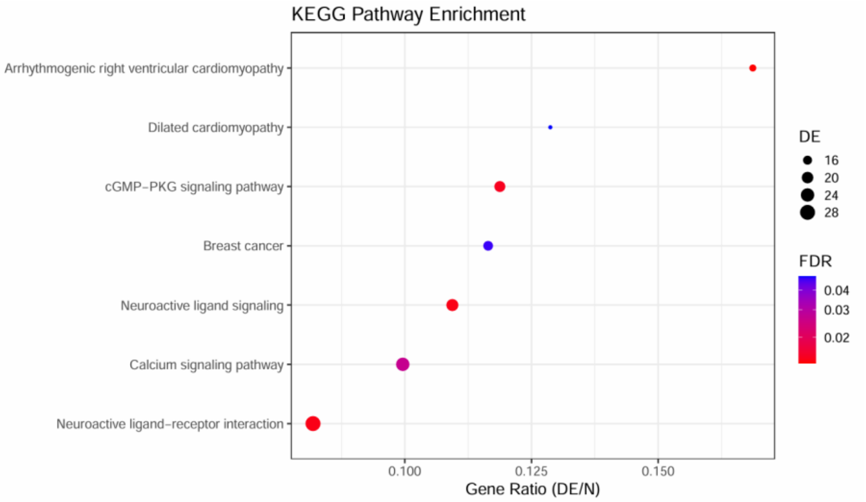


**Supplementary Fig. 7.** Gene Ontology (GO) term and KEGG pathway enrichment analysis. Dot plot showing enrichment of differentially methylated genes between *MLH1*-hypermethylated (n =42) and non-*MLH1*-hypermethylated (n =217) colorectal cancer samples, using missMethyl. The top 15 significantly enriched GO terms and pathways (FDR < 0.05) are shown. The gene ratio represents the number of differentially methylated genes (DE) in the term divided by the total number of genes in the term (N). The size of each dot corresponds to the number of DE genes in the term, and the colour indicates the false discovery rate (FDR).

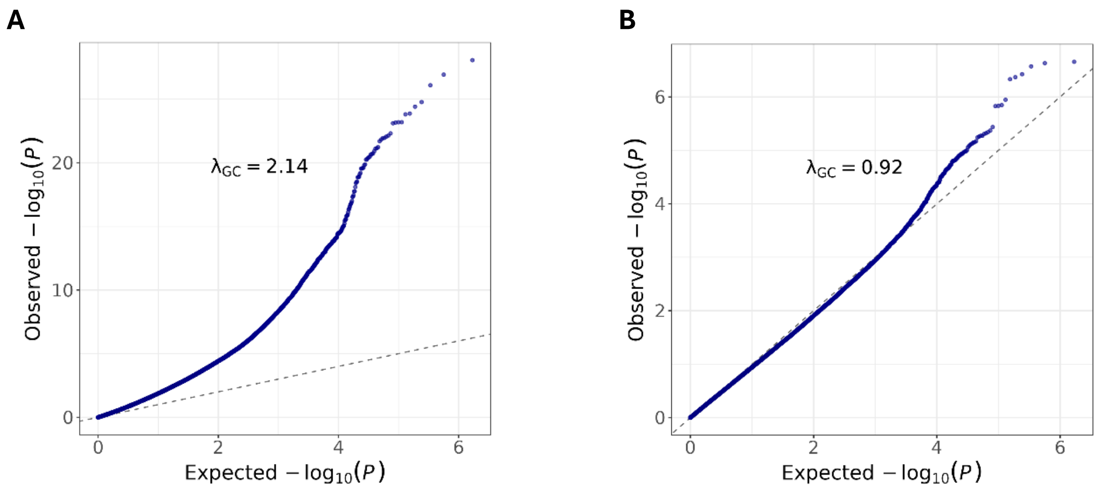


**Supplementary Fig. 8**. Quantile-quantile (QQ) plots of p-value distributions for EWAS analyses of the influence of clinical variables on methylation in colorectal cancer. Observed versus expected $-\log_{10}$ p-values for the EWAS of (A) anatomical location (right-sided vs left-sided), adjusted for MSI status, tumour purity, CIMP status and *MLH1* promoter methylation (B) patient age at diagnosis, adjusted for MSI status, tumour purity, anatomical location, CIMP status and *MLH1* promoter methylation. Lambda (λ) indicates the genomic inflation factor.

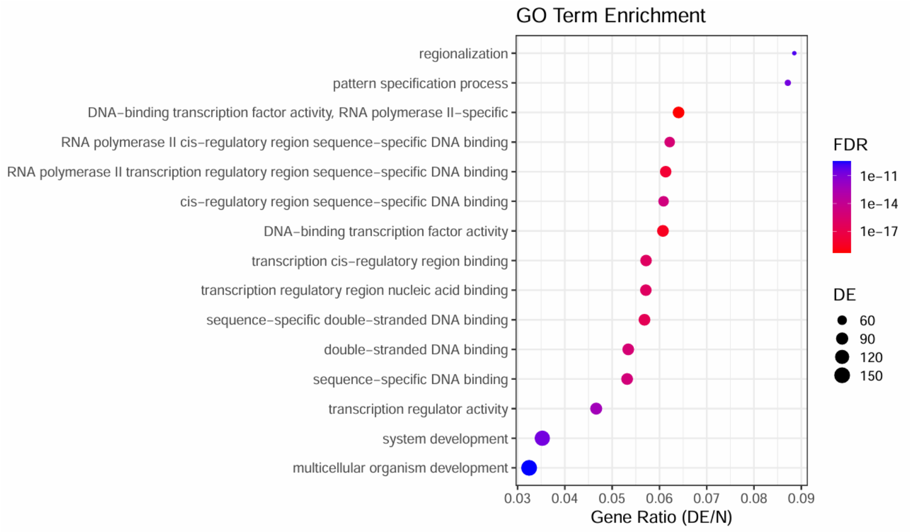


**Supplementary Fig. 9.** Gene Ontology (GO) term enrichment analysis. Dot plot showing enrichment of differentially methylated genes between left-sided (n =139) and right-sided (n =120) colorectal cancer samples, using missMethyl. The top 15 significantly enriched GO terms and pathways (FDR < 0.05) are shown. The gene ratio represents the number of differentially methylated genes in the term divided by the total number of genes in the term. The size of each dot corresponds to the number of differentially methylated genes (DE) in the term, and the colour indicates the false discovery rate (FDR). No KEGG pathways reached statistical significance.

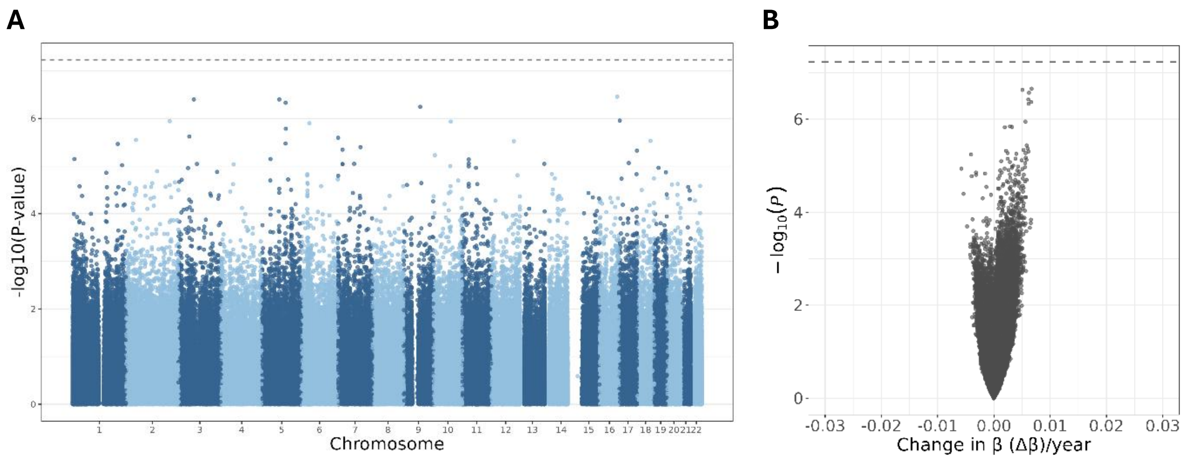


**Supplementary Fig. 10**. Epigenome-wide association study of patient age at diagnosis in colorectal cancer. (A) Manhattan plot showing $-\log_{10}$ p-values for CpG sites associated with patient age at diagnosis (n =259), adjusted for MSI status, tumour purity, anatomical location, CIMP status and *MLH1* promoter methylation. Bonferroni-corrected epigenome-wide significance ($p < 5.9 \times 10^{-8}$) is indicated by the grey dashed line (B) Volcano plot showing the association between methylation changes (Δβ per year of age) and statistical significance ($-\log_{10}$ p-value) for each CpG site, with the most significant genes labelled.

**A**

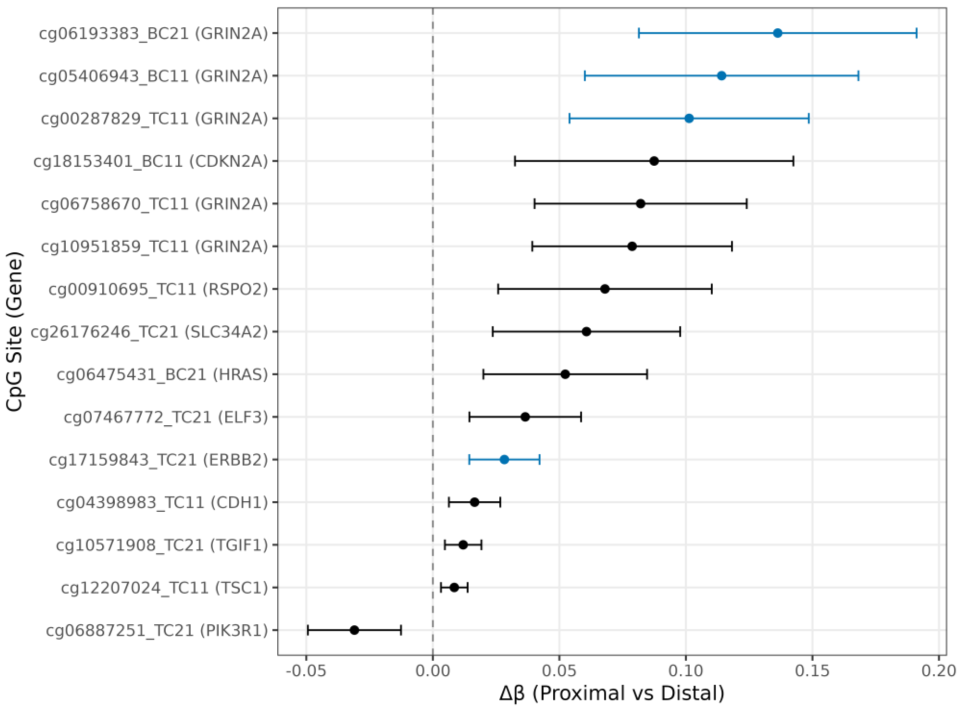


**B**

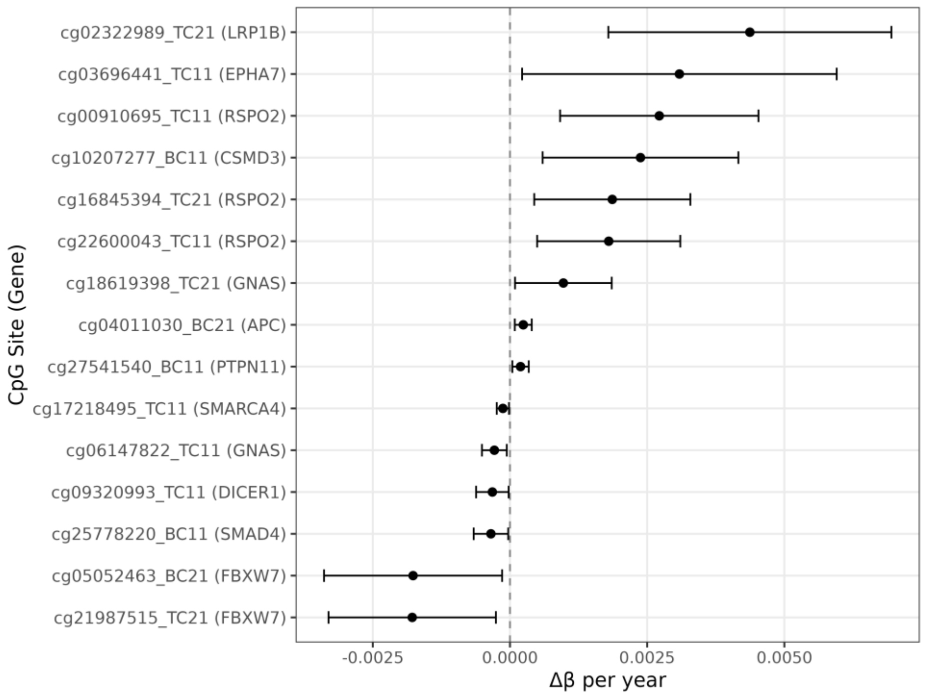


**Supplementary Fig. 11.** Targeted analysis of DNA methylation at CRC driver gene promoters. Forest plots showing associations between CpG methylation within the TSS200 regions of 82 driver genes and (A) anatomical location (proximal, n=120 vs distal, n=139), adjusted for MSI status, tumour purity, CIMP status, and *MLH1* promoter methylation or (B) age (n=259), adjusted for MSI status, tumour purity, anatomical location, CIMP status, and *MLH1* promoter methylation. Points represent estimated changes in methylation (Δβ) derived from limma moderated t-statistics, and horizontal lines show 95% confidence intervals. The top 15 CpG sites ranked by significance (p-value) are shown. Probes reaching statistical significance after Bonferroni correction ($p<1.04 \times 10^{-4}$) are highlighted in blue.
